# ISPRED-SEQ: Deep neural networks and embeddings for predicting interaction sites in protein sequences

**DOI:** 10.1101/2022.10.24.513521

**Authors:** Matteo Manfredi, Castrense Savojardo, Pier Luigi Martelli, Rita Casadio

## Abstract

The knowledge of protein-protein interaction sites (PPIs) is crucial for protein functional annotation. Here we address the problem focusing on the prediction of putative PPIs having as input protein sequences. The problem is important given the huge volume of sequences compared to experimental and/or computed protein structures. Taking advantage of recently developed protein language models and Deep Neural networks here we describe ISPRED-SEQ, which overpasses state-of-the-art predictors addressing the same problem. ISPRED-SEQ is freely available for testing at https://ispredws.biocomp.unibo.it.

## Introduction

Proteins are key players in most biological processes. Proteins are social entities and interact with membranes, within themselves or with other proteins, and/or biomolecules (including nucleic acids) to accomplish their functions within the cell. Among all the different features that protein functional annotation requires, it is also important to determine the likelihood of protein-protein interaction. Therefore, effective computational tools for the prediction of protein-protein interactions are important to characterize protein function and to further expand interactomes of different species [1–3].

The identification of Protein-Protein Interaction (PPI) sites, namely protein residues involved in physical interactions with protein partners, can be addressed using two complementary approaches. On one hand, different biochemical and biophysical experimental methods (such as X-ray crystallography, Nuclear Magnetic Resonance (NMR), alanine scanning mutagenesis and chemical cross-linking, to cite some) can be applied to determine protein-protein interfaces at the atomic or residue level [4]. Although very accurate, the applicability of these methods to large-scale characterization of PPI is still hampered by economical and technical issues.

On the other hand, computational methods are cost-effective solutions to complement experimental approaches in identifying and characterizing PPI sites. Docking programs are the major class of computational tools to study protein-protein interactions [for review, see ref 2]. Very accurate models can be obtained through docking when the two interacting partners are known in advance.

However, when the interacting partner/s is/are not known, machine-learning approaches can compute PPI sites on unbound protein chains. Historically, these methods have been relying on several physicochemical features extracted from protein sequence and/or structure and discriminate between interacting and non-interacting residues [2]..

The most accurate approaches are based on information extracted from protein 3D structures, including protein solvent accessibility, protrusion and depth indexes, secondary structures, B-factors, and others, including geometrical features, have been proven to be very informative [5]. Prediction of PPI sites from protein sequence alone is still challenging and methods developed for this specific task are less performing than the previous ones. Methods implemented so far for PPI prediction from protein sequence include in input evolutionary information, conservation scores and physical-chemical properties of amino acids (e.g., hydrophobicity, polarity, charge and/or conformational propensities). Eventually, structural features computed from protein sequence with specific classifiers, such as predicted solvent accessibility and secondary structure, are also included with the aim of filling the scoring gap with structure-based approaches. Several methods have been developed in the past and recent years [2], based mainly on different types of machine learning, including shallow and deep neural networks [6–15].

Recently, protein language models trained on large volumes of sequence datasets have been proven to be effective in providing protein/residue representations that are alternative and competitive with canonical hand-crafted features such as evolutionary information and physicochemical properties [16–19]. Representations/embeddings provided by these models have been successfully adopted in many prediction tasks [20–24].

Here we present ISPRED-SEQ, a novel webserver based on a deep learning model to predict PPI-sites from protein sequence encoded with an embedding procedure. The method stands on a deep architecture combining convolutional blocks and three cascading fully connected layers. ISPRED-SEQ is trained on a dataset of 6,066 protein chains derived from a dataset available in literature [14]. The main novelty of ISPRED-SEQ is the input generation, obtained using two state-of-the-art protein language models, ESM1-b [16] and ProtT5 [17].

We benchmark ISPRED-SEQ on four different independent test data derived from literature [9,14,15,25,26]. All proteins included in the training dataset have less than 25% sequence similarity with sequences in the testing sets, adopting a stringent homology-reduction procedure. Results show that ISPRED-SEQ performs at the state-of-the-art, reporting MCC scores higher than those obtained by other approaches in all the benchmarks performed.

The ISPRED-SEQ web server is freely accessible at https://ispredws.biocomp.unibo.it.

## Materials and Methods

### Datasets

#### Training dataset

For training the ISPRED-SEQ network we used a set of protein chains derived from a dataset available in literature [27] and already adopted, after some filtering steps, to train the DELPHI method [14]. The DELPHI dataset comprised 9,982 protein chain sequences extracted from the PDB and sharing no more than 25% pairwise sequence identity. Moreover, the sequences in the training set are also non-redundant (25% identity) with respect to all the sequences included in the independent test datasets (see next section). Starting from this set, we further restricted the number of protein sequences by filtering out all the chains (as in the correspondent UniProt file) having a coverage with the associated PDB structure/s less than 80%, in order to validate PPI annotation on structural experimental evidence. After this filtering step, we ended up with 6,066 protein sequences comprising 1,757,296 residues.

Annotation of PPI sites was then retrieved from the original data available from [27] and manually curated. Starting from the PDB structure of the complex, a residue of a given chain is defined in interaction if the distance between an atom of the residue and an atom of another residue in a different chain is below a given distance threshold, which routinely is set equal to the total sum of the van der Waals’ radii of the two atoms plus 0.5 Angstroms [27]. PPI annotations are available for the complete UniProt protein sequences after combining all interaction sites obtained from multiple protein complexes in which each protein is represented, adopting SIFTS [28] for the relative mapping of PDB and UniProt [27]. Overall, our dataset comprises 285,751 interaction sites, corresponding to about 16% of whole set of residues.

We split the training dataset into 10 different subsets for performing the 10-fold cross validation procedure. Before splitting, we further clustered the sequences at 25% sequence identity and 40% alignment coverage using MMseqs2 [29]. The cross-validation split was then performed by randomly distributing complete clusters (instead of individual sequences) among the different subsets. This step is required to capture residual local redundancies between pair of sequences that could have survived the first redundancy reduction performed during dataset construction.

#### Independent test datasets

To evaluate generalization performance of ISPRED-SEQ and to compare it with other state-of-the-art approaches we used four different independent test sets widely used in literature for comparative evaluation of tools [9,14,15,25,26]. Table 1 provide an overview of all datasets used in this study.

**Table 1.** Datasets

| Dataset | # Sequences | # Residues | # Interaction sites | # Non-interaction sites |
| --- | --- | --- | --- | --- |
| Training | 6,066 | 1,757,296 | 285,751 | 1,471,545 |
| Dset448<sup>†</sup> | 448 | 116,500 | 15,810 | 100,690 |
| Dset335<sup>†</sup> | 355 | 95,940 | 11,467 | 84,473 |
| Homo_TE | 95 | 24,766 | 5,564 | 19,202 |
| Hetero_TE | 48 | 14,056 | 1,313 | 12,743 |

<sup>†</sup>Both datasets include 34% and 36% of heterocomplexes, respectively

The first dataset comprises 448 protein chains used in a review comparing different tools for protein interaction site prediction from sequence [26]. The aim of the authors was to collect data including not only protein-protein interaction sites, but also annotations for DNA, RNA and small-ligand binding sites. For this reason, the dataset was obtained starting from the BioLip database [30], collecting nucleic-acid and ligand binding site annotations. For the set proteins retrieved from BioLip, authors also extracted protein-protein interaction sites by analyzing corresponding protein complexes available at the PDB. Protein interaction sites are identified using the same definition adopted for the training set (see above). Internal redundancy of the dataset was set to 25% pairwise sequence identity using the Blastclust tool [31]. We refer to this dataset to as the Dset448. The second dataset used here is referred to as the Dset335 and it is a subset of the Dset448 introduced in [14] for sake of comparing the methods DELPHI and DLPred [32]. The 335 sequences included in the dataset are indeed selected such that they are non-redundant at 25% sequence identity with the DLPred training set, hence enabling a fair comparison with this method. We used Dset335 to also include DLPred in our benchmark.

The third and fourth datasets, referred to as Homo TE and HeteroTE, respectively, were introduced by Hou and coauthors [9,25]. Recently, these sets were also used for evaluating the performance of the PIPENN prediction tool [15]. HomoTE and HeteroTE include 479 and 48 protein chains from homomeric and heteromeric complexes, respectively. Interface residues are defined in HomoTE and HeteroTE using a slightly different definition based on the computation of Accessible Surface Area (ASA) before and after complex formation: interacting residues are those whose ASA value undergo a change upon complex formation [25]. Nevertheless, as highlighted in literature [33], this definition provides very similar or equal interaction interfaces as those based on inter-chain distances.

### ISPRED-SEQ implementation

The ISPRED-SEQ general architecture is depicted in Figure 1. Staring from a protein sequence, ISPRED-SEQ input is constructed using two alternative protein language models: i) ESM1-b [16], an encoder-only transformer model trained on about 27 million sequences from UniRef50 [34], and ii) ProtT5 [17], a sequence-to-sequence model derived from the T5 architecture [35], trained on the large Big Fantastic Database (BFD) [36] comprising 2.1 billion sequences and fine-tuned on the UniRef50 database.

**Figure 1.**
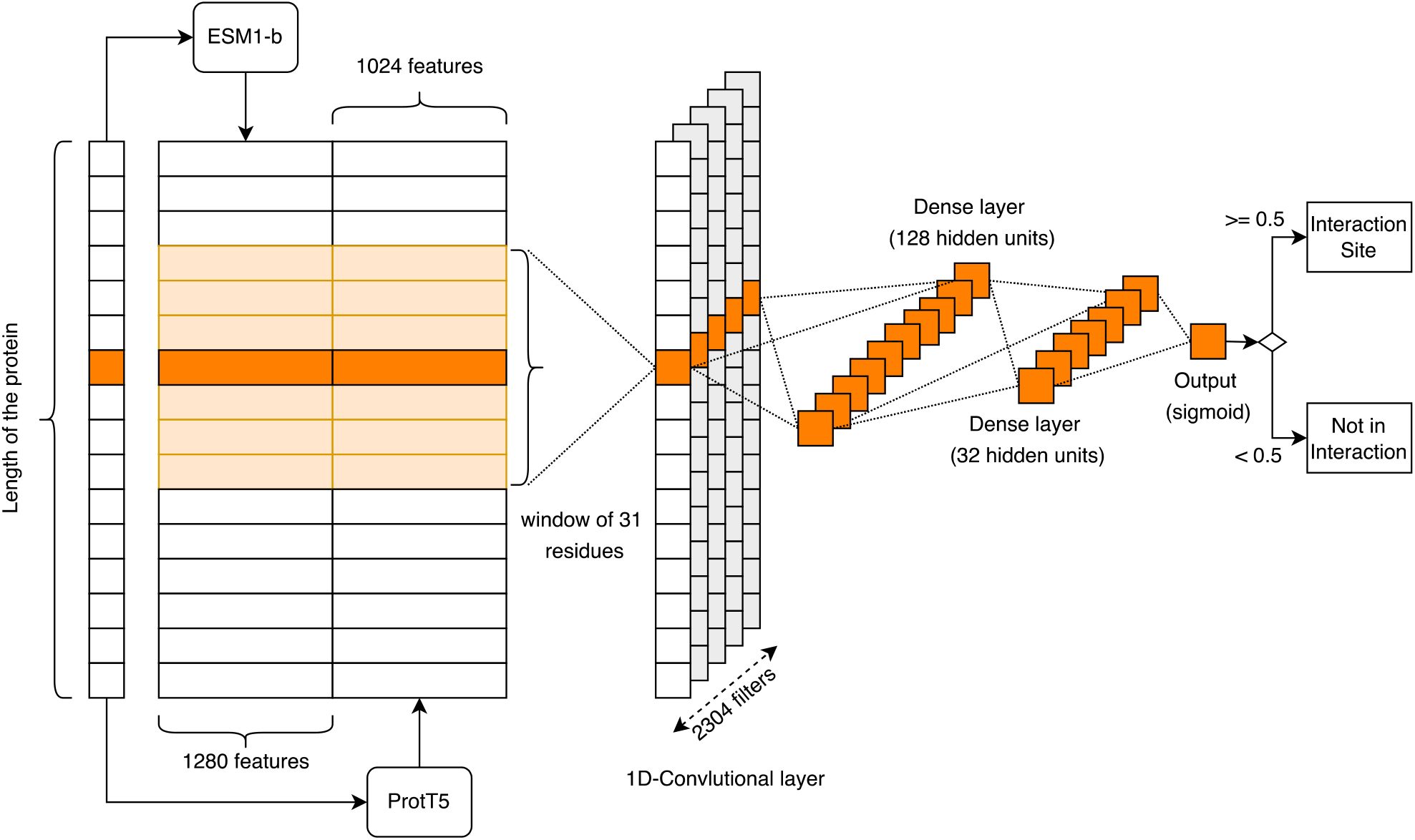
The ISPRED-SEQ deep network architecture

For each residue in the input sequence, ESM1-b and ProtT5 provide embeddings of dimension 1280 and 1024, respectively. These are then concatenated to form a single vector comprising 2304 components for each residue.

Since ESM1-b can only accept in input sequences of length lower than 1022, all longer sequences are split into non-overlapping chunks of equal length such that none is longer than 1022. After this step, the sequence embedding is reconstructed by concatenating all the chunks.

The joint embedding (ESM1-b + ProtT5) is then processed using a four-layer network. The first layer is a 1-dimensional convolutional neural network with 2304 filters (the number of filters is set as to be equal to the input dimension) and a filter width of 31, corresponding to a window comprising 31 flanking residues and centered at each residue position. The positional output of the convolutional layers is processed by two dense, fully connected layers with 128 and 32 hidden units, respectively. The final output consists of a single unit with a sigmoid activation function. Each residue is classified as interaction site if the output value is greater or equal to 0.5, as not in interaction otherwise.

For sake of assessing the contribution of the input encoding, we also trained alternative models based on different types of inputs, including: the sequence one-hot encoding, providing 20 values per residue, the position-specific scoring matrix (PSSM), computed using two runs of HHblits [37] against the UniClust30 database [38] and providing 20 values per residues, ESM1-b embedding only (1280 values per residue) and ProtT5 embedding only (1024 values per residue). For all the models trained, we adopted the same architecture shown in Figure 1, and changing the number of convolutional filters to be equal to the input dimension (20 for one-hot and PSSMs, 1280 for ESM1-b and 1024 for ProtT5).

Training is performed using minibatches of 64 residues adopting an early stopping procedure that halts the training after 10 epochs without a decrease in the validation loss. The loss that we implemented is a binary cross-entropy and we adopted an Adam optimizer [39].

To fix all the hyperparameters of the model we performed a grid search using a strict 10-fold cross validation. After that, we retrained the final model on the whole training dataset, and we evaluated it on the different benchmark sets.

### Scoring measures

The following measures were used to score performance of the different methods:

- Accuracy (Q2):

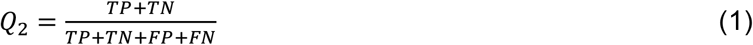
- Precision:

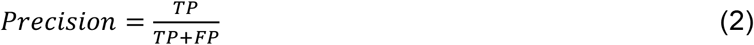
- Recall:

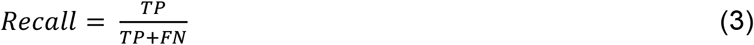
- F1-score, the harmonic mean of precision and recall:

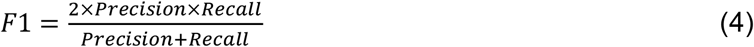
- Area Under the Receiver Operating Characteristic Curve (ROC-AUC).
- Matthews Correlation Coefficient (MCC):

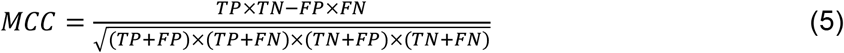

Routinely, the probability value discriminating between positive and negative predictions is set to 0.5. For benchmarking on blind test sets ISPRED-SEQ with other approaches [14,15,26], we adopted a methodological strategy previously described [26]. According to this procedure, for each of the different methods, a method specific threshold is introduced to set the number of positive predictions equal to the number of real positive examples [14,15,26]. AUC values are however independent of the threshold values.

## Results

### ISPRED-SEQ performance

For fine tuning ISPRED-SEQ, we tested the network architecture using a 10-fold cross-validation procedure to compare different input encodings. Specifically, we evaluated five different models trained on different inputs, including: i) the sequence one-hot encoding, ii) the sequence profile, iii) the ESM1-b embedding only, iv) the ProtT5 embedding only and v) the joint embedding obtained combining ESM1-b and ProtT5. Table 2 lists the results.

**Table 2.** ISPRED-SEQ 10-fold cross-validation results using different input encodings

| Input | MCC | F1 | Precision | Recall | Q2 | AUC |
| --- | --- | --- | --- | --- | --- | --- |
| One-hot | 0.08±0.01 | 0.28±0.01 | 0.20±0.01 | 0.49±0.06 | 0.60±0.04 | 0.58±0.01 |
| Profiles | 0.14±0.01 | 0.32±0.01 | 0.20±0.01 | 0.79±0.04 | 0.46±0.04 | 0.64±0.01 |
| ESM1-b [16] | 0.30±0.01 | 0.42±0.01 | 0.30±0.01 | 0.70±0.03 | 0.68±0.02 | 0.74±0.01 |
| ProtT5 [17] | 0.31±0.01 | 0.43±0.01 | 0.31±0.01 | 0.72±0.02 | 0.69±0.01 | 0.75±0.02 |
| ESM1-b+ProtT5 | 0.34±0.01 | 0.45±0.01 | 0.32±0.01 | 0.75±0.05 | 0.70±0.02 | 0.80±0.02 |

Models incorporating canonical features (one-hot and sequence profiles) are both outperformed by embedding-based approaches. MCCs obtained with embedding-based approaches score with values above 0.30 and higher that the 0.14 value obtained with only the sequence profile as input (Table 2). Data are shown in Table 2, obtained adopting a cross validation procedure. This highlights the effectiveness of language model representations in the task of predicting PPI sites. The two different language models (ESM1-b and ProtT5) provide similar contributions individually achieving comparable MCC scores (0.30 and 0.31, respectively). When combined, the value of MCC is 0.34, suggesting that the ESM1-b and ProtT5 are complementary, and their combination is advantageous for the problem at hand. This configuration is our final ISPRED-SEQ.

To compare ISPRED-SEQ performance with other state-of-the-art tools, we adopted the same strategy described in [14,15] and defined in [26] by which binary predictions are obtained using a different threshold for each method such that the number of positive predictions (FP+TP) is equal to the number of real positive examples (TP+FN). State-of-the-art tools include: DELPHI [14], PIPENN [15], SCRIBER [11], SSRWF [8], CRFPPI [40] and LORIS [41]. Table 3 shows the results.

**Table 3.** Comparative benchmark on different independent test sets

| Method | Dataset | MCC | F1 | Precision | Recall | Q2 | AUC |
| --- | --- | --- | --- | --- | --- | --- | --- |
| ISPRED-SEQ (th=0.5) | Dset448 | 0.34 | 0.42 | 0.29 | 0.78 | 0.71 | 0.82 |
| ISPRED-SEQ (th⇒FP=FN) | Dset448 | 0.39 | 0.47 | 0.47 | 0.47 | 0.86 | 0.82 |
| DELPHI [14]<sup>†</sup> | Dset448 | 0.27 | 0.37 | 0.37 | 0.37 | 0.83 | 0.74 |
| PIPENN [15]<sup>‡</sup> | Dset448 | 0.25 | 0.39 | 0.39 | 0.39 | 0.79 | 0.73 |
| SCRIBER [11]<sup>†</sup> | Dset448 | 0.23 | 0.33 | 0.33 | 0.33 | 0.82 | 0.72 |
| SSWRF [8]<sup>†</sup> | Dset448 | 0.18 | 0.29 | 0.29 | 0.29 | 0.81 | 0.69 |
| CRFPPI [40]<sup>†</sup> | Dset448 | 0.15 | 0.27 | 0.26 | 0.27 | 0.81 | 0.68 |
| LORIS [41]<sup>†</sup> | Dset448 | 0.15 | 0.27 | 0.26 | 0.26 | 0.81 | 0.66 |
| ISPRED-SEQ (th=0.5) | Dset335 | 0.33 | 0.40 | 0.27 | 0.77 | 0.72 | 0.82 |
| ISPRED-SEQ (th⇒FP=FN) | Dset335 | 0.39 | 0.46 | 0.46 | 0.46 | 0.87 | 0.82 |
| DELPHI [14]<sup>†</sup> | Dset335 | 0.28 | 0.36 | 0.36 | 0.36 | 0.85 | 0.75 |
| SCRIBER [11]<sup>†</sup> | Dset335 | 0.23 | 0.32 | 0.32 | 0.32 | 0.84 | 0.72 |
| DLPred [32]<sup>†</sup> | Dset335 | 0.21 | 0.31 | 0.31 | 0.31 | 0.84 | 0.72 |
| ISPRED-SEQ (th=0.5) | Homo_TE | 0.42 | 0.56 | 0.42 | 0.83 | 0.71 | 0.84 |
| ISPRED-SEQ (th⇒FP=FN) | Homo_TE | 0.46 | 0.58 | 0.58 | 0.58 | 0.81 | 0.84 |
| PIPENN [15]<sup>‡</sup> | Homo_TE | 0.34 | 0.49 | 0.49 | 0.49 | 0.77 | 0.77 |
| ISPRED-SEQ (th=0.5) | Hetero_TE | 0.20 | 0.27 | 0.17 | 0.68 | 0.65 | 0.72 |
| ISPRED-SEQ (th⇒FP=FN) | Hetero_TE | 0.16 | 0.24 | 0.24 | 0.24 | 0.86 | 0.72 |
| PIPENN [15]<sup>‡</sup> | Hetero_TE | 0.11 | 0.20 | 0.20 | 0.20 | 0.85 | 0.66 |

<sup>†</sup> Data taken from [14]. <sup>‡</sup>Data taken from [15].

Performance of all methods, with the exclusion of ISPRED-SEQ, are extracted from literature [14,15]. Specifically, performance on Dset448 and Dset335 for DELPHI, SCRIBER, SSRWF, CRFPPI and LORIS are derived from [14], while results of PIPENN in all datasets are taken from the original reference paper [15]. All the methods tested provide numerical prediction scores representing the propensity of each input residue to be a PPI site. For our ISPRED-SEQ, performance measures obtained using this strategy are labelled as “th⇒FP=FN” in Table 3. A direct comparison with the state-of the art methods is therefore possible. For sake of completeness, we also show ISPRED-SEQ score obtained using the threshold of 0.5 on the output prediction score. This threshold assumes a probability meaning for the output of ISPRED-SEQ and it is the one adopted in the web server.

Regardless of the method adopted for choosing the threshold, Table 3 indicates that ISPRED-SEQ outperforms all the methods in all the considered datasets. In the Dset448 (the most recent and complete dataset released in literature so far [26]), ISPRED-SEQ achieves a MCC value of 0.39, twelve percentage points higher than the one obtained by the second top-performing method, DELPHI.

In the Homo-TE dataset containing homomeric interfaces, ISPRED-SEQ reaches a MCC value of 0.46, again significantly higher than the one registered by PIPENN. Performance on the small Hetero-TE are lower. However, also in this case, ISPRED-SEQ outperforms the other tested method (PIPENN) by several percentage points.

Independently of the procedure adopted for evaluating the scoring indexes, ISPRED-SEQ overpasses the performance of all other methods. This is also evident when considering the AUC values reported in Table1, totally independent of the strategy adopted for the other scoring indexes.

## Conclusions

In this paper we present ISPRED-SEQ, a novel method for the prediction of PPI sites from sequence. ISPRED-SEQ novelty is the adoption of input encodings based on embeddings generated by two state-of-the-art protein language models, ESM1-b and ProtT5. In our tests, residue representations based on embeddings outperform canonical feature descriptors such as one-hot encoding and sequence profiles.

We evaluated ISPRED-SEQ using several independent datasets released in literature and compared its performances against recently state-of-the-art approaches also based on deep-learning algorithms. In all the tests performed, ISPRED-SEQ significantly outperformed top-scoring methods, reaching MCC scores of 0.39 on recent benchmark datasets containing more than 300 proteins.

We propose ISPRED-SEQ as a valuable tool for the characterization of protein interface residues starting from the protein primary sequence.

We released ISPRED-SEQ as a publicly accessible web server available at https://ispredws.biocomp.unibo.it.

## Acknowledgments

The work was supported by PRIN2017 grant (project 2017483NH8_002), delivered to CS by the Italian Ministry of University and Research. We acknowledge ELIXIR-IIB, the Italian node of the ELIXIR infrastructure.

